# Making waves: Comparative analysis of gene drive spread characteristics in a continuous space model

**DOI:** 10.1101/2022.11.01.514650

**Authors:** Mingzuyu Pan, Jackson Champer

## Abstract

With their ability to rapidly increase in frequency, gene drives can be used to modify or suppress target populations after an initial release of drive-containing individuals. Recent advances in this field have revealed many possibilities for different types of drives, and several of these have been realized in experimental demonstrations. These drives all have unique advantages and disadvantages related to their ease of construction, confinement, and capacity to act as a modification or suppression system. While many properties of these drives have been explored in modelling studies, assessment of these drives in continuous space environments has been limited, often focusing on outcomes rather than fundamental properties. Here, we conduct a comparative analysis of many different gene drive types that have the capacity to form a wave of advance against wild-type alleles in one-dimensional continuous space. We evaluate the drive wave speed as a function of drive performance and ecological parameters, which reveals substantial differences between drive performance in panmictic versus spatial environments. In particular, we find that suppression drive waves are uniquely vulnerable to fitness costs and undesired CRISPR cleavage activity that can form resistance alleles in embryos by maternal deposition. Some drives, though, retain robust characteristics even with widely varying performance characteristics. To gain a better understanding of drive waves, we compare panmictic performance of drives across the full range of drive frequencies. We find that rates of wild-type allele removal in panmictic setting is correlated with drive wave speed, though this is also affected by a range of other factors. Overall, our results provide a useful resource for understanding the performance of drives in continuous spatial environments, which may be most representative of potential drive deployment in many relevant scenarios.

## Introduction

From a modest start, the field of gene drive has grown rapidly in the past 20 years, especially after the advent of CRISPR/Cas9-based drive systems^1–3^. Gene drives are alleles that have the potential ability to spread through a whole population by biasing inheritance in their favor. This could make them powerful tools for controlling vector-borne diseases, invasive species, and other pests. Thus far, different types of gene drives have been demonstrated in several species, including yeast^4,5^, fruit flies^6–10^, mosquitoes^11–14^, and mice^15^, with many additional gene drives having been modeled in various settings^1–3^.

These gene drives can be divided into categories in multiple possible ways. Some gene drives are designed to modify populations, which could prevent insects from transmitting diseases such as malaria and dengue. Other drives are designed to suppress populations, which could be used to directly eliminate pest species. Besides application, gene drives can be separated into classes based on how confined they would be to a specific target population. When the introduction frequency of a drive is higher than the introduction threshold, the gene drive can spread rapidly. If the introduction frequency is lower than the threshold, it’s expected that the drive allele will be lost from the population. Zero-threshold drives such as CRISPR homing drives and Y-linked X-shredders are fast, powerful drives that can be considered “global” drive systems due to their ability to invade from small starting populations^16–21^ (though high fitness costs can sometimes move these into more confined categories). An intermediate category consists of “regional” drives that lack an introduction threshold when they have ideal performance but gain one if there are any imperfections in the drive that lead to even a small fitness cost (and occasionally other types of imperfections for some drive types). Examples of this sort of system include *Medea*^22^ and TARE^23–25^/ClvR^26,27^ drives. For underdominance alleles, drive or wild-type homozygotes are more reproductively successful than drive/wild-type heterozygotes. One feature of this kind of gene drive is its non-zero threshold under all conditions, making them “local” drives systems that are highly confined but spread more slowly^28–36^.

Properties of these gene drives have been extensively explored with computational modeling. However, most of these models simulate panmictic populations or linked, discrete populations connected with fixed migration^21,37,46,47,38–45^. While such models can be important for gaining a fundamental understanding of gene drive behavior and would apply well to certain situations, many realistic settings would instead involve spatially continuous populations. In such settings, a species could potentially move over a wide range, but individuals would generally spend their life in only a small part of this range, mating and producing offspring in only their local area.

When a gene drive spreads in such settings, it tends to form a “wave of advance” as it moves into a wild-type population.

Some efforts have been made to understand fundamental drive properties in spatially explicit settings using dense networks of linked populations^48–51^ or abstract patches^52^, though it is unclear how such results would apply to spatially continuous setting. Other studies have assessed the characteristics of drive wave advance into a population of wild-type individuals for Driving Y/X-shredders^53,54^, homing drives^55–58^, and underdominance systems^59–61^. For underdominance drives, some insights can also be gained by general analytical assessments^62,63^. Together, these represent a substantial body of knowledge, yet there are major gaps in our fundamental understanding of spatial gene drive behavior. Many newer drives have not been assessed in these models. Other models use drive mechanisms that have not been observed in real drives, such as zygotic conversion of wild-type alleles to drive alleles by homing drives (only resistance alleles have been observed to form in zygote/early embryo stages, with drive conversion taking place in germline cells). Many more recent studies of gene drive in continuous space have focused on outcomes in complicated situations, such as the chasing phenomenon for suppression drives^54,64– 70^, rather than fundamental spatial properties. Thus, there exists a need to reassess the spatial performance of both new and older drive designs using a common framework to facilitate comparisons between drive types under different conditions.

In this study, we investigated the ability of several different drives to spread in continuous spatial environments. We measured the wave speed these drives, comprehensively covering several major and minor variants with different performance and ecological parameters. We also assessed the qualitative and quantitative manner in which these drives increase in panmictic populations and how this related to their spatial wave speed. Together, these results can help us predict and better understand the critical features of how drives behave in realistic, spatially continuous environments.

## Methods

### Panmictic Model

Our simulations start from a panmictic population model in SLiM software (version 3.8)^71^. We used discrete, non-overlapping generations, and each generation begins with reproduction and ends with removal of adults from the previous generation. Initially, every fertile female will sample a male from the whole population. When the male is sampled, the probability of him becoming the mate is proportional to his fitness, which is based on his genotype. If the male she chooses is sterile, then she will not reproduce. This is representative of mosquito populations where females typically only mate once^72,73^.

In this panmictic population model, when a female selects a mate successfully, we define her fecundity as 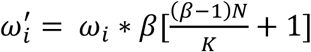. In this equation, *N* is the population size, *β* is the low-density growth rate in this population, *K* is the carrying capacity, and *ω*_*i*_ is the female’s fitness, which is based on her genotype. To determine the actual number of offspring produced, we draw from a binomial distribution where we use *n* = 50 samples and set the probability of creating an offspring with each sample as 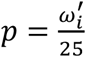. This represents 50 eggs with independent survival. The number 50 is chosen because it is likely close to the upper limit of the number of offspring a successful female could have under permissive conditions for most species (actual numbers of eggs could be greater, but this usually occurs when offspring mortality rates are high, even in good environments). This allows the possible number of offspring to have a reasonably sized distribution when the low-density growth rate is at its maximum and density competition is low. When the population size is close to its capacity, fecundity will converge to 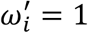. This will result in two offspring for each female on average, but if the population size is lower than capacity, each female will average more than two offspring. We set the low-density growth rate to 6 by default in this model, which is based on estimates of the *Anopheles* mosquito rate of between 2 and 12^44^. Offspring receive a genotype based that takes drive activity into account.

Nonviable offspring are immediately removed. See the results section for details of how each drive functions.

To get the genotype-based fitness, we usually use multiplicative drive allele fitness, assuming that each allele has an independent fitness cost. For individuals that are drive homozygotes, individuals with only one chromosome where the drive can reside (such as males with drives on the Y chromosome), or *Wolbachia*-infected individuals, their “genotype”-based fitness is equal to the drive fitness value (set to 1.0 by default, which is equal to wild-type fitness), while for individuals that are drive heterozygotes, their genotype-based fitness is equal to the square root of drive fitness. For 2-locus drives, we assume that only the first drive allele subtype has the cargo gene and thus a fitness cost. For some types of suppression drives that do not carry cargo, we consider different fitness costs (see drive classification section for details). For somatic fitness costs, we apply the fitness value only to drive/wild-type heterozygotes. In most cases, such fitness costs are only applied to one sex.

Our simulations were initialized by allowing an initial wild-type population of *K* = 100,000 individuals to equilibrate for 10 generations. For gene drives that have a high introduction threshold, including 2LTADDE, 2LTADE, 2LTARE, TAHRE, and *Wolbachia*, we then released drive homozygous individuals. For other gene drives, we released drive heterozygotes. For the Driving Y/X-shredder and the Z-linked W-shredder, we only released males with one drive allele into to the population, and for the X-linked Y-shredder, we released only female heterozygotes. We released males and females at equal proportions for all other gene drive types.

### Determination of integrals from panmictic drive simulations

To better understand how each drive removed wild-type alleles from the population (which could simply be removed or replaced by drive alleles), we obtained a standardized integral. For each panmictic model, we collected the number of wild-type alleles and the wild-type allele frequency of the whole population for each generation, combining several simulations with different starting frequencies with ten replicates each. Then, for each generation transition, we obtained the absolute fractional change of the number of wild-type alleles (relative to the starting number of alleles in the population, which was 200,000) by subtracting the number of wild-type alleles in the current generation from the number of wild-type alleles in the next generation and dividing these results by 200.000. Using these data, we the obtained the integral of the absolute fractional change of the number of wild-type alleles over the full range of wild-type allele frequency from 0 to 1.

### Spatial Model

In the spatial model, we extended our panmictic framework into one-dimensional space with a total length of 1. Reproduction is the same as in the panmictic model, except that some events have a spatial range. In particular, females will only sample males from a restricted circle rather than the whole population, with a radius equal to the *migration value*. If the female can’t find a male after the maximum number of attempts, she will not reproduce.

Local competition also affects the fitness of each female and is responsible for density regulation in the model. We define an expected carrying density strength, *ρ*_*k*_ *=* 1/2 * *K* * *interaction distance*, and then we define a local density *ρ*_*i*_. This is the local density strength for each female within the *interaction distance* around her, which we set to 0.01. Each individual contributes a density strength of “1” if they are at the same position as the focal female, and this strength linearly declines to zero as the distance increases to 0.01 (thus, individuals contribute an average of 1/2 density strength in our one-dimension space, accounting for the factor of 1/2 in our equation for *ρ*_*k*_). The fitness of the female is then defined as 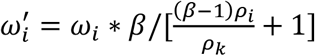, where other parameters are the same as the panmictic model. Offspring are placed according to a normal distribution with a standard deviation equal to the *migration value*, which is set to 0.04 by default. If an offspring’s location falls outside of our space, they are assigned a new position until it falls within the boundaries.

Our one-dimensional spatial simulations were initialized by randomly scattering a population of *K=*10,000 wild-type individuals into the whole space. Then, we let the population equilibrate for 10 generations. After that, we released drive individuals into the left edge of the arena, with the exact release pattern dependent on the drive and sufficient to ensure that the drive will establish a wave if this is possible.

### Determination of Wave Speed

We divided the whole one-dimensional space into 10 equal slices. In our simulations, we collected the gene drive frequency in each slice at each generation. To measure the drive wave speed, we wanted to ensure that the wave had enough time to reach its advancing equilibrium state, and we wanted to end the simulation before the wave would be affected by the edge of the arena. We thus chose slices on the position scale between 0.2 and 0.3 and between 0.7 and 0.8. When the gene drive frequency in the first slice was more than 0.5 for the first time, we defined this generation as *S*_1_ as “start generation” and the frequency as *G*_1_ as “start frequency”. The generation before the start generation is defined as *S*_0_, with a drive frequency of *G*_0_. Then we calculated the start generation *Start* using the following equation:

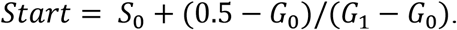

This approximation reduces error from using discrete generations and is designed to find a hypothetical continuous generation number where the drive frequency would be 0.5.

We then repeated this process for the second slice to find the *T*_1_, “stop generation”, the previous

*T*_0_ generation, and the final frequency in these generations, *F*_1_ and *F*_0_, respectively. We then similarly calculated the stop generation *Stop* by

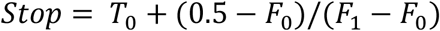

For some gene drives, the equilibrium level of the drive allele frequency (after it has increased to its maximum) in a panmictic population was sometimes below 0.5 or too close to this, resulting in inaccurate measurements. These included the both-sex lethal, both-sex sterile, female-sterile, and haplodiploid homing suppression drives. For these drives, we set the trigger point for measurements to be the first generation in which the wild-type allele frequency fell below 0.8, and we similarly collected the *start* and *stop* generations based on this level.

The distance between the center of the first and second slices is 0.5, so we can thus estimate the wave speed for all of the drives as:

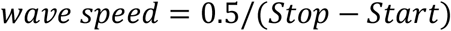

### Data generation and software

Simulations were carried on the High-Performance Computing Platform of the Center of Life Science at Peking University. We used Python and R to analyze the data and draw figures. We also used Origin Lab to calculate integrals. All SLiM models, data, and scripts are available on Github (https://github.com/jchamper/ChamperLab/tree/main/Drive-Wave-Speed).

## Results

### Classification of drive mechanisms in this study

The gene drives that we consider in this study (Table 1) represent most major designs for self-sustaining systems that are capable of forming a wave of advance against wild-type alleles, at least under some circumstances. While we do not consider all possible interesting drive types that have been imagined thus far (such as binary expression drives^46^, double drives^74^, HD-ClvR^75^, t-haplotypes drives^76^, *Medusa*^77^, *Semele*^78^, or other toxin-antidote variants^79^), we still assess most demonstrated types and many other recent designs. These can be broadly divided into several categories.

**Table 1.**
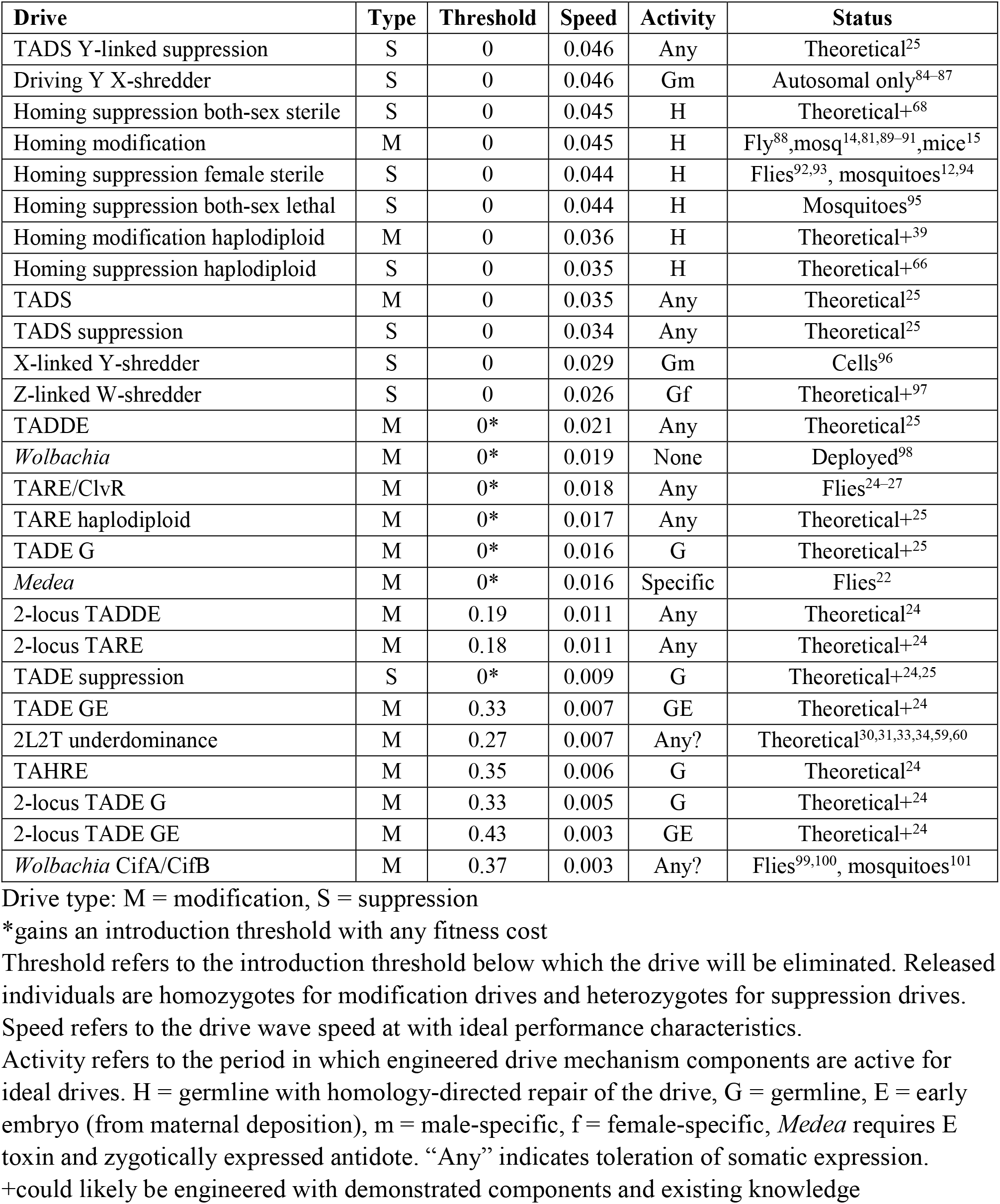
List of gene drives in this study.

Most well-known are the homing drives. These use a nuclease, usually CRISPR/Cas9, to cleave a wild-type allele in the germline. Homology-directed repair then leads to copying of the drive allele into the DNA break, thus converting the wild-type allele to a drive allele. The rate that this happens is represented by the drive efficiency. Remaining wild-type alleles can be converted to resistance alleles in the germline. We assume that multiplex gRNAs are used, so all resistance alleles are nonfunctional, which is less determinantal to drives than functional resistance alleles^47^. We also assume that this occurs in half of remaining wild-type alleles. Additionally, wild-type alleles can be converted to resistance alleles in the early embryo from due to maternal deposition of Cas9 and gRNA if the mother has at least one drive allele. We model several different types of homing drives with different target genes. For modification drives, we assume that the drive provides rescue, so only resistance alleles are dominant-lethal for haplolethal target genes and recessive-lethal for haplosufficient but essential target genes. The latter of these would tend to functional similarly to the prominent class of homing drives called “integral drives”^80,81^ if the target gene met preferred functional requirements (essential but haplosufficient, and fully functional in drive alleles). For suppression drives, genotypes that lack any wild-type alleles will suffer depending on the target, which could result in female sterility, both-sex sterility, or both-sex lethality. Fitness effects for these drives are somatic, applying only to drive/wild-type individuals and only to females for female fertility genes. We chose somatic fitness costs in these cases because it is much more likely that the drive will suffer from these (suppression drives lack cargo but potentially have some limited leaky somatic Cas9 expression) than from standard fitness costs based around number of cargo genes in the genotype. We also model homing drives in haplodiploid organisms, where males form from unfertilized eggs and can thus not have any drive activity because they only have one of each chromosome. X-linked drives would have the same population dynamics^39^. For haplodiploid modification rescue drive, we only consider haplosufficient but essential targets. For haplodiploid suppression drive, we only consider female fertility targets because other targeting strategies would not allow the drive to be successful^66^.

Toxin-Antidote Dominant Sperm (TADS) drive is another form of CRISPR drive, but it does not copy itself by homology-directed repair. Instead, the germline efficiency is simply the germline cut rate. This drive targets a gene that is expressed after meiosis I and is required for spermatogenesis. The drive provides a rescue copy. Thus, sperm with a disrupted target gene are non-viable unless the sperm also carries a drive allele. We model TADS drive alleles as located in the target gene (“same-site”) for the modification form. Same-site TADS suppression drive targets a male fertility gene with additional gRNAs (with the same cut rates as the main drive) while sitting at the TADS target site. We thus model it with male-specific somatic fitness costs. “Distant site” TADS suppression sits in the male fertility gene and targets only the TADS target gene. We model it with normal fitness costs.

Another class of gene drives induces population suppression by biasing the sex ratio, resulting in population suppression due to scarcity of one sex. The classic example of this is the driving Y/X-shredder, which sites on the Y chromosome and eliminates the X-chromosome at a rate equal to the drive efficiency. A normal number of offspring are produced, but these have a biased sex ratio due to elimination of female gametes. A homing drive that contains an X-shredder^82^ would tend to act like a driving Y/X-shredder if its components are highly efficient. TADS Y-linked suppression drive functions similarly, but with a different mechanism as described above. The drive efficiency refers to the germline cleavage rate of TADS target alleles (assumed to be on an autosome) in the presence of the drive. Both of these drives would be difficult to engineer due to difficulties in knocking transgenes into the Y-chromosome and then achieving sufficiently high expression. Another method for biasing the sex ratio is to place a Y-shredder on the X-chromosome, the reducing the frequency of males in the population. In ZW chromosome systems (where males are ZZ, and females are ZW, with W having similar characteristics to Y chromosomes), Z-linked W-shredders would effectively reduce the number of females based on the shredding efficiency. We assume that maternal deposition of nuclease does not occur in ZW females with the drive, though this could modestly increase drive performance by eliminating some females that escape germline W-shredding activity.

All of the previously described gene drives are “zero-threshold” systems, though some will gain thresholds if fitness costs are very high. Another class of gene drives lack a threshold in ideal form but gain one with any fitness costs. Several members of this family are CRISPR toxin-antidote drives, where drive efficiency refers to the germline cut rate of the target gene (embryo cleavage is also possible and has a variable effect on these drives). All cutting forms nonfunctional resistance alleles, sometimes known as “disrupted” alleles for these non-homing drives, and drive alleles have a recoded rescue site that cannot be cut. The most prominent CRISPR toxin-antidote drive is Toxin-Antidote Recessive Embryo (TARE), often known as Cleave and Rescue (ClvR) in its “distant site” form, where the drive is not located in the same gene as its target. These drives provide rescue for a haplosufficient but essential target gene, so individuals with two disrupted alleles are nonviable at the embryo stage (unless a drive is present elsewhere in the genome for the distant-site form, which we do not model here but has generally similar performance^25^).

Several variants exist for TARE drive with similar threshold properties. One is a haplodiploid version of TARE. In another the target site is haplolethal, so disrupted alleles are dominant-lethal, and the drive is known as Toxin-Antidote Dominant Embryo (TADE). Because TADE is a more powerful drive, it can be used for population suppression. In same-site TADE suppression, the drive sites in its haplolethal target and additionally targets a female-fertility drive with extra gRNAs. This drive has female somatic fitness costs. We also model distant site TADE suppression (normal fitness costs), which sites inside the female fertility gene and only has gRNAs for the haplolethal target. Another variant in this class is Toxin-Antidote Double-rescue Dominant Embryo (TADDE). This drive is the same as TADE except that the rescue is doubled, so drive/disrupted allele heterozygotes are still viable.

*Medea* drive is based on RNAi and follows simple rules. If an offspring lacks a drive allele and has a drive heterozygous mother, it will be nonviable at a rate equal to the drive efficiency. *Medea* has similar properties to TARE drive.

We also model a non-drive system based on *Wolbachia* bacteria, which can spread through a population in a similar manner to a gene drive. Females infected with *Wolbachia* pass it on to all their offspring, but *Wolbachia*-infected males cannot pass on *Wolbachia*. However, wild-type females that mate with *Wolbachia* males do not have any offspring due to cytoplasmic incompatibility. *Wolbachia* gains an introduction threshold if it has any fitness costs.

The final class of gene drives consists of those that have an introduction threshold even in ideal form. Most of these have underdominance properties, where drive heterozygotes are less fit than either drive or wild-type homozygotes. These include TADE drive with embryo cutting, as well as another form of CRISPR toxin-antidote drive called Toxin-Antidote Half-rescue Recessive

Embryo (TAHRE). TAHRE functions similarly to a TARE drive, but two copies of the drive are required for the rescue element to be sufficient, so drive/disrupted allele heterozygotes are nonviable. TAHRE functions best without embryo cutting, but its threshold only increases from 35% to 41% with 100% embryo cutting due to maternally deposited Cas9 and gRNA.

One class of CRISPR underdominance gene drives are 2-locus systems. These are similar to the basic forms, except that there are two subtypes of the drive, with each subtype being located at a separate locus. Each of these drives targets the gene that the other provides rescue for (we assume same-site rescue for all 2-locus drives considered here), so many genotypes without at least one of each drive type tend to be nonviable (though ultimately this depends on cleavage rates of the target genes). We consider 2-locus variants with TARE, TADE, and TADDE drives and targets (including TADE drives with only germline “G” cutting or with both germline and embryo “GE” cutting). Aside from CRISPR systems, a well-known 2-locus, 2-toxin-antidote (2L2T) underdominance system is considered in which any drive-carrying individuals are only viable if they have at least one copy of each drive subtype.

The final drive system considered is one based off of the *Wolbachia* mechanism, but instead of actual *Wolbachia*, this drive contains the *Wolbachia* phage genes CifA and CifB that are responsible for *Wolbachia’s* cytoplasmic incompatibility^83^. Inheritance in this drive is normal, except when a wild-type female mates with a male containing at least one copy of the CifA/CifB drive allele. In this case, the female’s fecundity is reduced by the drive efficiency parameter.

Self-limiting drives, which are designed to eventually be removed from a population, can form a wave of advance in some situations, but such waves would never reach an equilibrium state because they would quickly break down due to their temporary nature. We thus do not consider such drives in this study. We also do not consider weaker gene drives that cannot advance against a wild-type population. Though still potentially useful in a variety of situations, these drives would require widespread releases in spatial environments and would be unable to form waves of advance without a major geometric advantage^59^. These tend to be drives with an introduction threshold of equal to or greater than 50% (see Table S1 for a list of prominent, self-sustaining drives that could not form a wave of advance). Finally, we do not explicitly model tethered drives^24,37^, though the individual components of tethered drives would behave nearly identically to other drives that we model.

### Qualitative characteristics of drive waves

To visualize advancing drive waves, we set up “snapshots” of ideal drives that had advanced approximately halfway through the arena (Figure 1, S1). For suppression drives (the Driving Y and homing suppression drives in Figure 1, plus several drives in Figure S1), snapshots showed that they eliminated most of the population in the left side of the arena successfully. However, the X-linked Y-shredder (Figure S1) failed to suppress the population when the drive wave had advanced halfway through because its female-bias actually creates more potential reproduction in our model (where males can mate an unlimited number of times). In the long term, the population would eventually be eliminated after males are gone, and elimination can occur sooner if males are limited in their mating capacity^96^. In all modification drives (such as the homing modification drive in Figure 1), the left half of the arena shows successful population modification and replacement of wild-type individuals, though most CRISPR toxin-antidote variants such as TARE in Figure 1, *Medea*, and *Wolbachia* CifA/CifB drives (Figure S1) take a long time for drive heterozygotes to be replaced by drive homozygotes. Note that TADE drives do not have this drawback (Figure S1).

**Figure 1.**
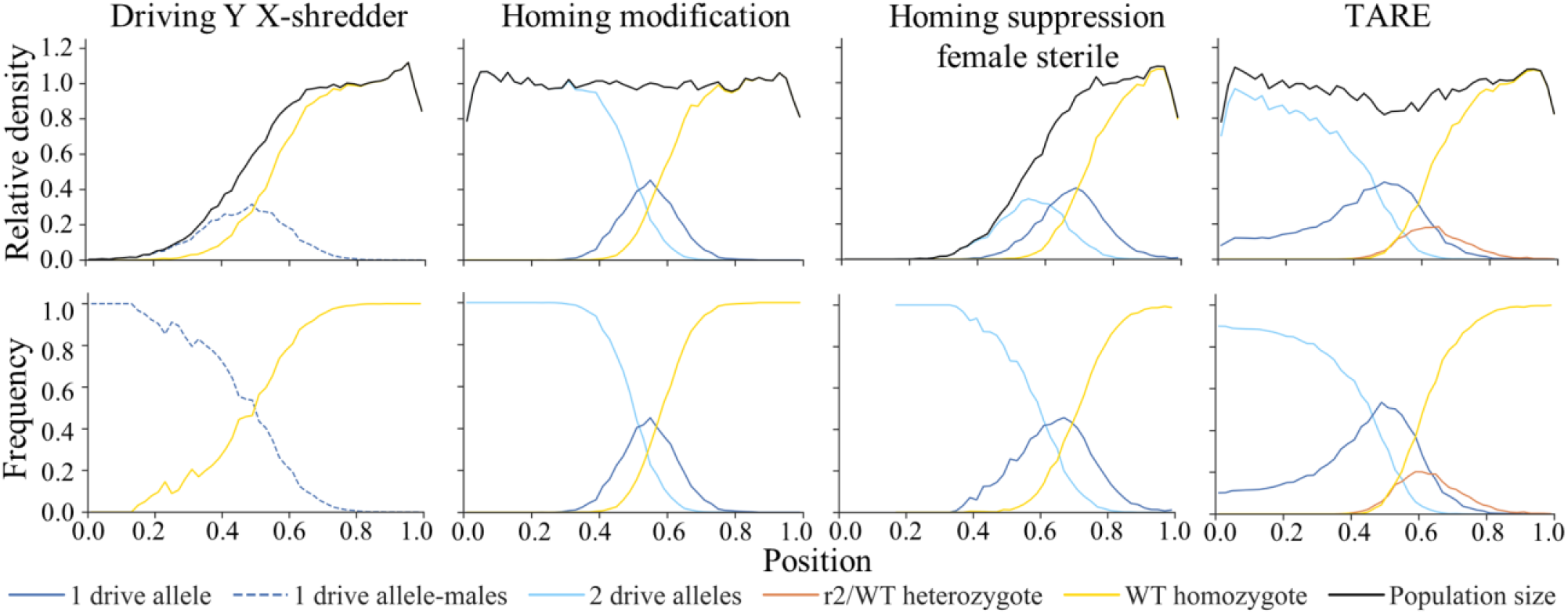
Advancing drive waves. The relative density (compared to wild-type in the absence of drive) of several genotype categories is displayed for an advancing drive wave, as well as the total population density. The lower panels show the frequencies of each genotype category. The graphs are divided into 50 points covering the whole one-dimensional space of length 1.

One prominent feature of many modification drives (Figure 1 TARE drive, several drives in Figure S1) is that the population density is lower where the wave front of the drive intersects with wild-type. This does not occur in homing drives, but in all the other drives, a toxin-antidote effects results in some offspring being nonviable. This reduces the population only where drive and wild-type individuals co-occur. Because TADS drive only cause sterility in males with two nonfunctional resistance alleles and no drives alleles, this population density reduction is much less intense than other drives. This sort of population reduction in any drive could reduce the speed of the advancing drive wave, because even though the drive is increasing in frequency in these regions, the reduced total population will reduce the number of drive individuals migrating into the wild-type region.

Some types of drives appear to have a particularly small or narrow advancing drive wave. This is best represented by TADE GE variants where many heterozygotes are eliminated due to nonfunctional resistance alleles (Figure S1). Such drives have difficulty moving forward because many drive alleles are eliminated at the wave front in addition to wild-type alleles. TADE suppression drive has a very wide wave front due to its suppressive effect and weaker ability to spread compared to other suppression drive systems (Figure S1).

### Drive wave speeds

We first measured drive wave speed under ideal drive performance characteristics (Table 1). A clear pattern emerged. The fastest drives were the zero-threshold Y-linked drives and homing drives because these were potentially capable of doubling in crosses between drive and wild-type individuals. Next were the haplodiploid homing drives and TADS drives, which can double, but only on females and males, respectively. Their speed in panmictic populations is half that of autosomal homing drives, but their wave speed was well over half the speed of homing drives, underscoring the difference between drive spread in panmictic versus spatial models. Next were other sex-linked shredders. These were substantially slower than Y-linked editors because they are often found in individuals without a target chromosome to shred, which contributes nothing toward increasing the drive frequency. The following group consisted of frequency dependent drives that lacked an introduction threshold in ideal form, but gain one if there are any fitness costs. These intermediate drives spanned a narrow range of speeds and included both TARE drive and *Wolbachia* bacteria. Notably missing from this group was TADE suppression drive, which was substantially slower due to its suppressive effect. Indeed, TADE suppression drive fell in the middle of the last group of drives, all the rest of which had introduction thresholds in ideal form. These had diverse mechanisms (such as 2-locus systems) and widely varying speeds, which loosely correlated with their introduction thresholds.

By varying the migration rate for several representative drives (Figure S2), we found that the wave advance velocity was always proportional to the migration rate, as expected.

### Assessment of drive wave speed based on drive efficiency

We next varied the germline drive efficiency (Figure 2), or for drives that can also have embryo cutting, both the germline efficiency and embryo cut rate (Figure 3, S3). Both of these parameters play an important role in the drive allele’s transmission process and should thus have a large impact on the wave advance speed. Migration and low-density growth rate were fixed at their default values, and the drives were assumed to have no fitness cost.

**Figure 2.**
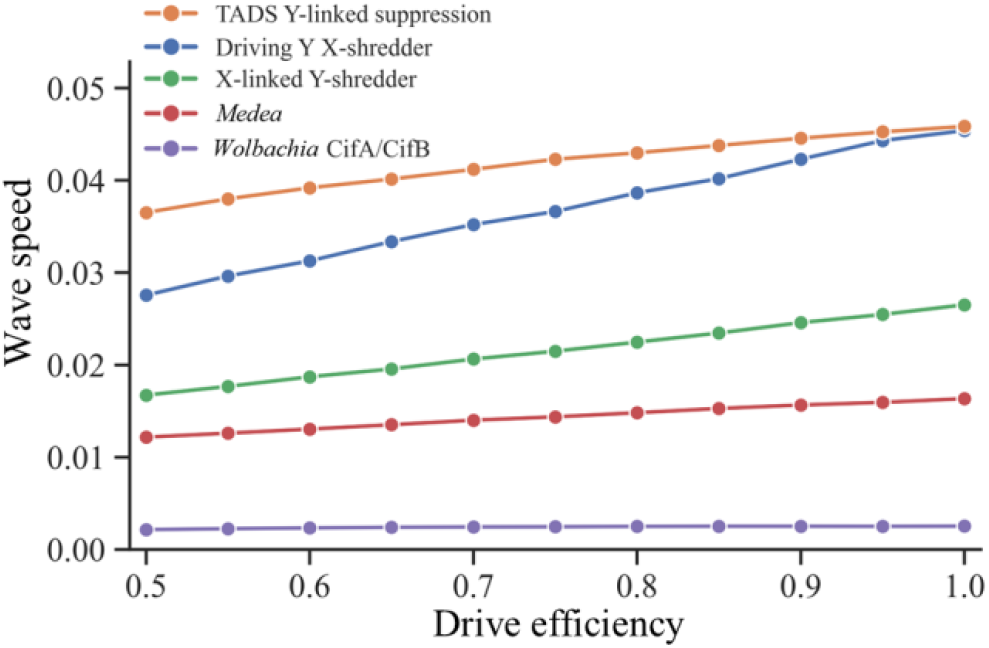
Line graph showing the effects of drive efficiency. Drive carriers were released into the left edge of a population of wild-type individuals, and the wave speed was measured for varying drive efficiency. Each point represents the average from at least 200 simulations.

**Figure 3.**
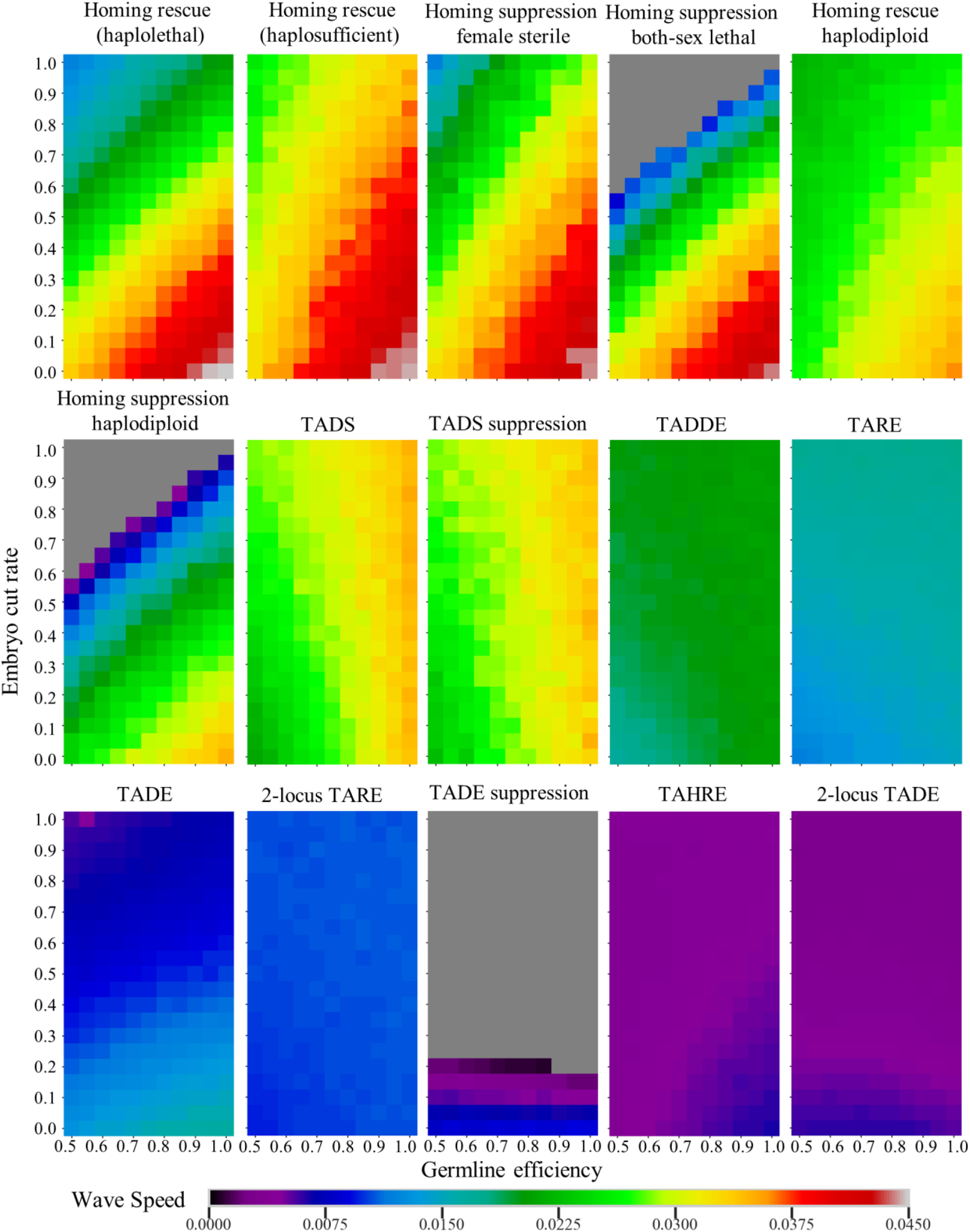
Effect of germline efficiency and embryo cut rate on wave speed. Drive carriers were released into the left edge of a population of wild-type individuals, and the wave speed was measured for varying germline efficiency and embryo cut rate. Each point represents the average of 20 simulations. Grey means that a wave of advance could not form.

In general, the germline efficiency will almost always help drive allele transmission. However, reducing this parameter to the minimum value we considered (0.5) had little negative effect on the frequency-dependent *Medea* and *Wolbachia* CifA/CifB drive (Figure 2), as well as the CRISPR toxin-antidote drives (Figure 3, S3). In some of these drives, embryo cutting could also “cover” for the reduced germline efficiency to restore drive wave speed. Interestingly, the optimal germline efficiency for 2-locus TADE value was well below 1, because this resulted in less removal of drive alleles in the presence of disrupted alleles.

Embryo cutting can sometimes help but it will usually hurt a gene drive, depending on the drive type (Figure 3, S3). This occurs when the transmission of the drive allele is undermined, and the wave speed is then slower. Homing drives and TADE drives are greatly slowed by embryo cutting in particular. For TADE drives, this is because embryo cutting removes large numbers of drive alleles which can alone greatly increase the threshold of these drives. For homing drives, this tends to be because resistance alleles prevent drive conversion (though for haplolethal rescue homing drive, drive alleles are also removed, hence the greater effect of embryo cutting compared to rescue homing drives targeting haplosufficient genes). Compared to modification drives, a higher embryo cut rate has more negative effect on suppression drive, even causing failure of the drive to form a wave in less powerful homing drive systems and especially in TADE suppression, which can only form a wave if the drive’s embryo cut rate is below ~0.2.

2-locus TADE and TAHRE are modestly slowed by embryo cutting, which tends to remove more drive alleles. TADDE was fully tolerant of embryo cutting because it causes no harm to the drive, but it also doesn’t help if there is high germline cleavage in males and females. 2-locus TARE and TADDE were also tolerant of embryo cutting because drive and wild-type alleles are removed in similar proportions. Embryo cutting was actually helpful for TADS drives and the single-locus TARE system because these drives can remove wild-type alleles faster with embryo cleavage without any removal of drive alleles. For TADS, though, the boost provided from embryo cutting was generally of small magnitude if germline efficiency was high because most drive increase comes from germline activity in males.

### Assessment of drive wave speed based on fitness costs and ecological parameters

We also measured the wave speed of gene drives with varied fitness value and low-density growth rate (Figure 4, S4), fixing germline efficiency and 1 and the embryo cut rate at either 0 or 1 (whichever was better for the drive). For drives that are more sensitive to fitness costs, results were plotted separately (Figure 5).

**Figure 4.**
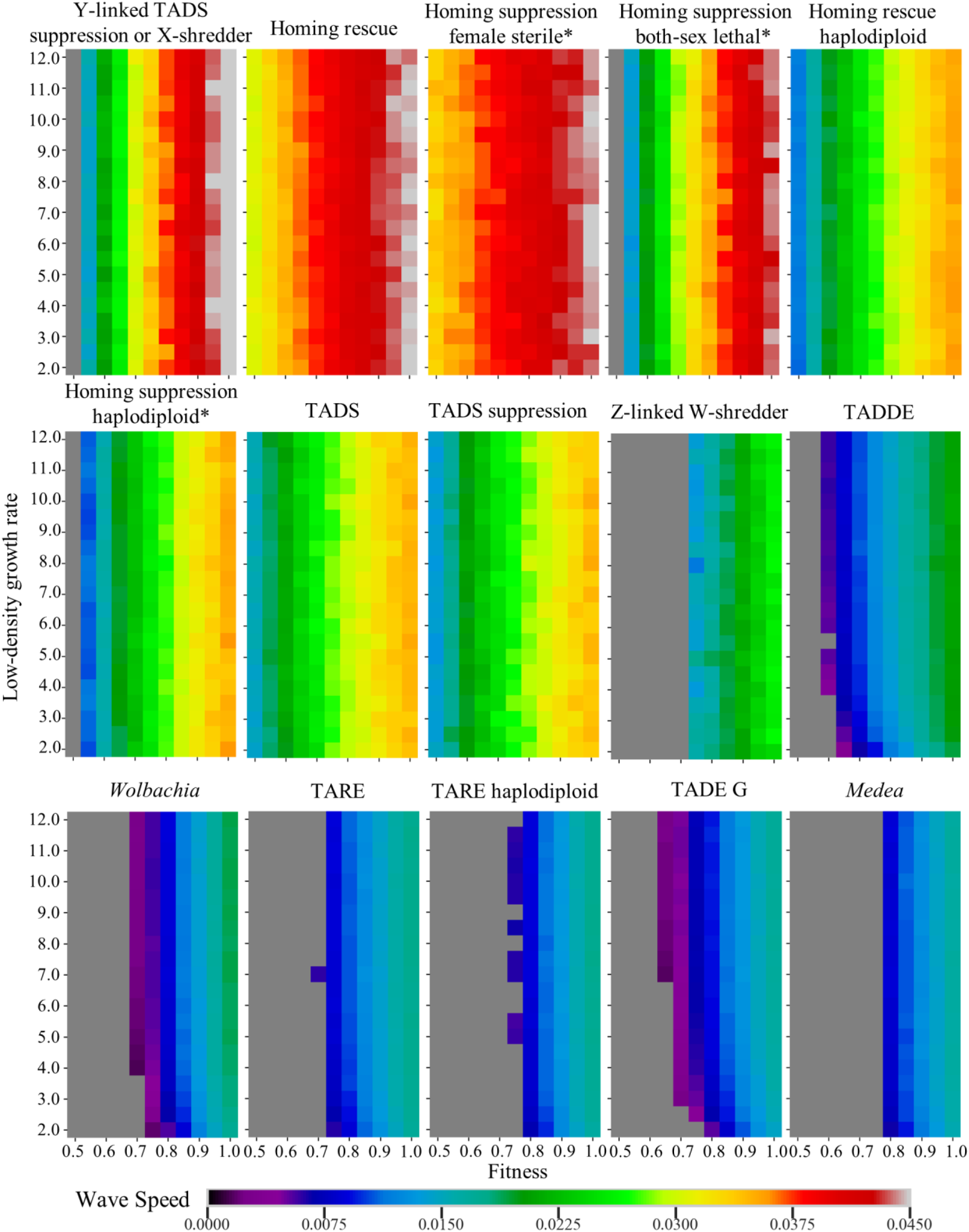
Effect of low-density growth rate and fitness value on wave speed. Drive carriers were released into the left edge of a population of wild-type individuals, and the wave speed was measured for varying fitness and low-density growth rate. Each point represents the average of 20 simulations. Grey means that a wave of advance could not form. * means that somatic fitness costs were used instead of multiplicative fitness.

**Figure 5.**
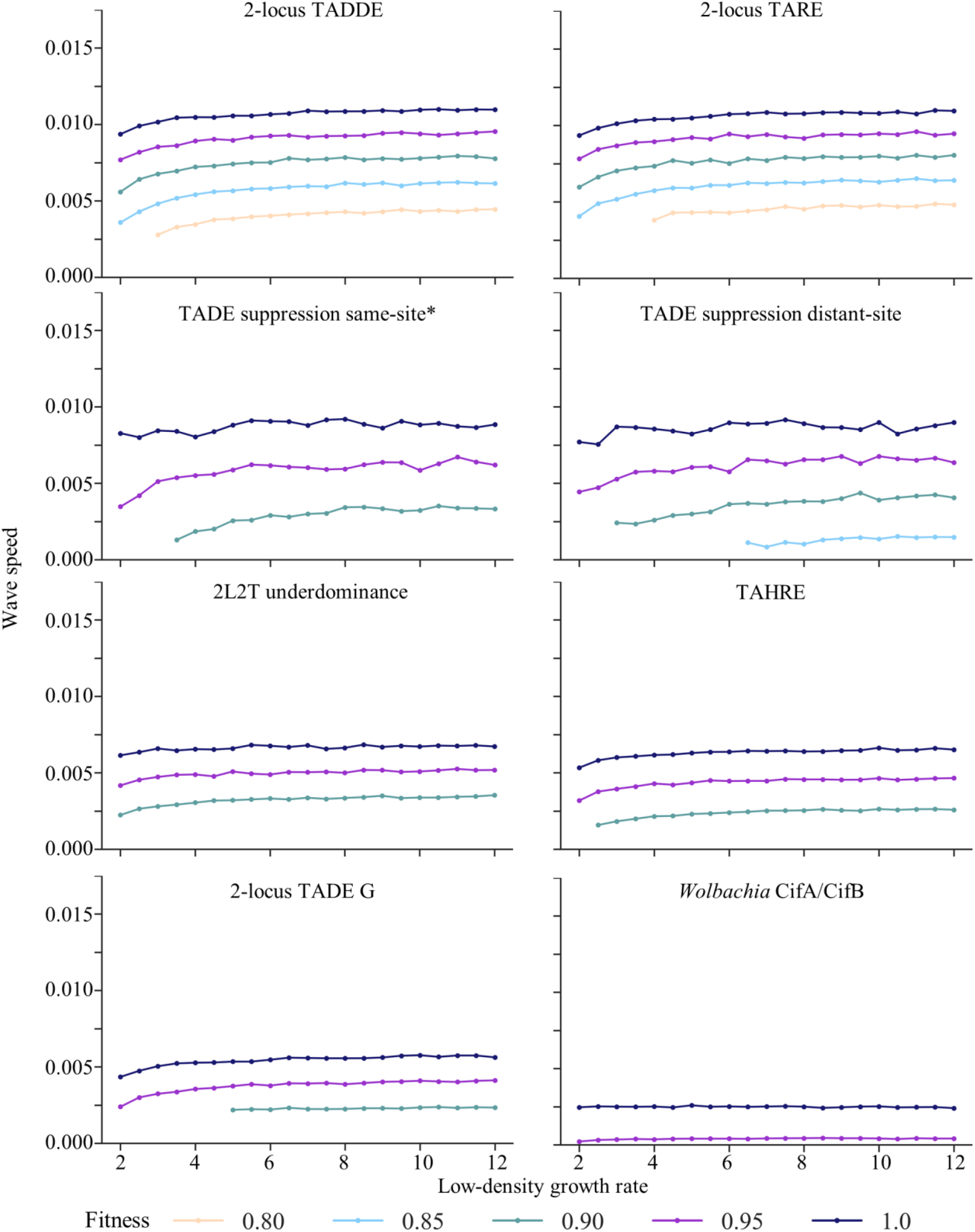
Effect of low-density growth rate and fitness value on wave speed for highly confined drives. Drive carriers were released into a population of wild-type individuals, and the wave speed was measured for varying fitness and low-density growth rate. Lower fitness values (including below 0.8) that don’t appear resulted in failure to form a wave. Each point represents the average of 20 simulations. * somatic fitness costs were used.

Fitness costs could be due to the drive itself, but if well-engineered, the primary fitness cost of a drive would be caused by its cargo. However, for suppression drives that target an essential gene without rescue, fitness costs can also commonly be caused by undesired somatic nuclease expression and cleavage. In both cases, fitness was in relation to wild-type (which was assumed to have a fitness of 1) and was assumed to proportionately affect male mating success and female fecundity. For all the drives we assessed, reductions in fitness caused a major decrease in the wave speed. For some drives, a sufficiently low fitness prevented the drive from being able to form a wave in the first place. This was seen in all the frequency-dependent drives, but also in the weaker suppression drives for the fitness range we considered (from 0.5 to 1). This was related to the high introduction threshold of these drives in the presence of fitness costs. When the threshold reaches 0.5, the drive has no advantage over wild-type alleles in well mixed populations and would thus not be expected to form a wave in which the drive would have higher frequency over about half the wave and lower frequency in the other half. In practice, drives would fail to advance a wave when the threshold was a little below 0.5 because the density of the population would be lower where the drive has high frequency, reducing migration of the drive into the wave front compared to migration of wild-type individuals. This feature could potentially be less pronounced in models with a different type of density dependence.

The low-density growth rate is the relative growth rate of the population when the density is low and competition becomes unimportant. Homing drives and other fast drives that do not make any individuals nonviable were unaffected by the low-density growth rate. However, higher values will result in reduced suppression from any drive effects, which can have a negative effect on the outcome of a suppression drive release^65,68,70^. On the other hand, this effect could potentially increase the number of drive individuals, especially in suppression drives, and thus increase the speed of the wave. In practice, though, increasing this parameter had little positive effect on any drive in terms of wave speed. Zero-threshold suppression drives would still not be heavily affected by this because the front of the wave is responsible for most of its advance, and this area does not experience substantial suppression. For many frequency-dependent drives with toxin-antidote, effects, however, a small boost in drive speed was seen between low-density growth rates of around 2-5. High low-density growth rates could also allow a wave to form in situations where fitness costs would otherwise make this difficult.

For comparison purposes, we also investigated the wave speed of wild-type individuals advancing into empty space (Figure S5), which can be important during chasing^54,65,66,68,70^. The wave speed of wild-type populations was quite a bit faster when the low-density growth rate was higher because this is the main mechanism by which the population will increase. In general, this was also faster than any drive by the time the low-density growth rate reached around 3, even compared to rapid, zero-threshold drive systems. However, increasing the low-density growth rate past 10 did not result in significant increases in wild-type wave speed because by this point, the migration/dispersal rate was the limiting factor (even small numbers of migrants to a new area could rapidly increase in density).

### Drive spread characteristics in panmictic populations

To better understand the relative performance of drives in panmictic populations, we analyzed the ability of ideal drives to increase in frequency across the full spectrum of possible frequencies. Rather than using a large number of starting frequencies, we initialized the simulation with a small number of starting frequencies near the drive threshold (on either side of the threshold for drives with nonzero thresholds). This would allow us to understand drive performance along its natural trajectories, rather than a series of artificially chosen points that may have different ratios of drive heterozygotes, homozygotes, and other genotypes.

Based on this data, we calculated the relative change of drive allele frequency for all the gene drives we considered (Figure 6). This was obtained for each transition between two generations by finding the difference in drive frequency between the generations divided by the frequency in the earlier generation. Negative values mean that the drive is decreasing in frequency (this usually only occurs when the drive frequency is below its introduction threshold, though note that the thresholds in Table 1 represent drive homozygous releases rather than a natural balance of drive homozygotes and heterozygous as seen in this data). These results can help show how the drive will perform across the whole wave of advance, though the “width” of the wave that corresponds to each frequency will be variable and can’t be directly inferred from this panmictic drive data.

**Figure 6.**
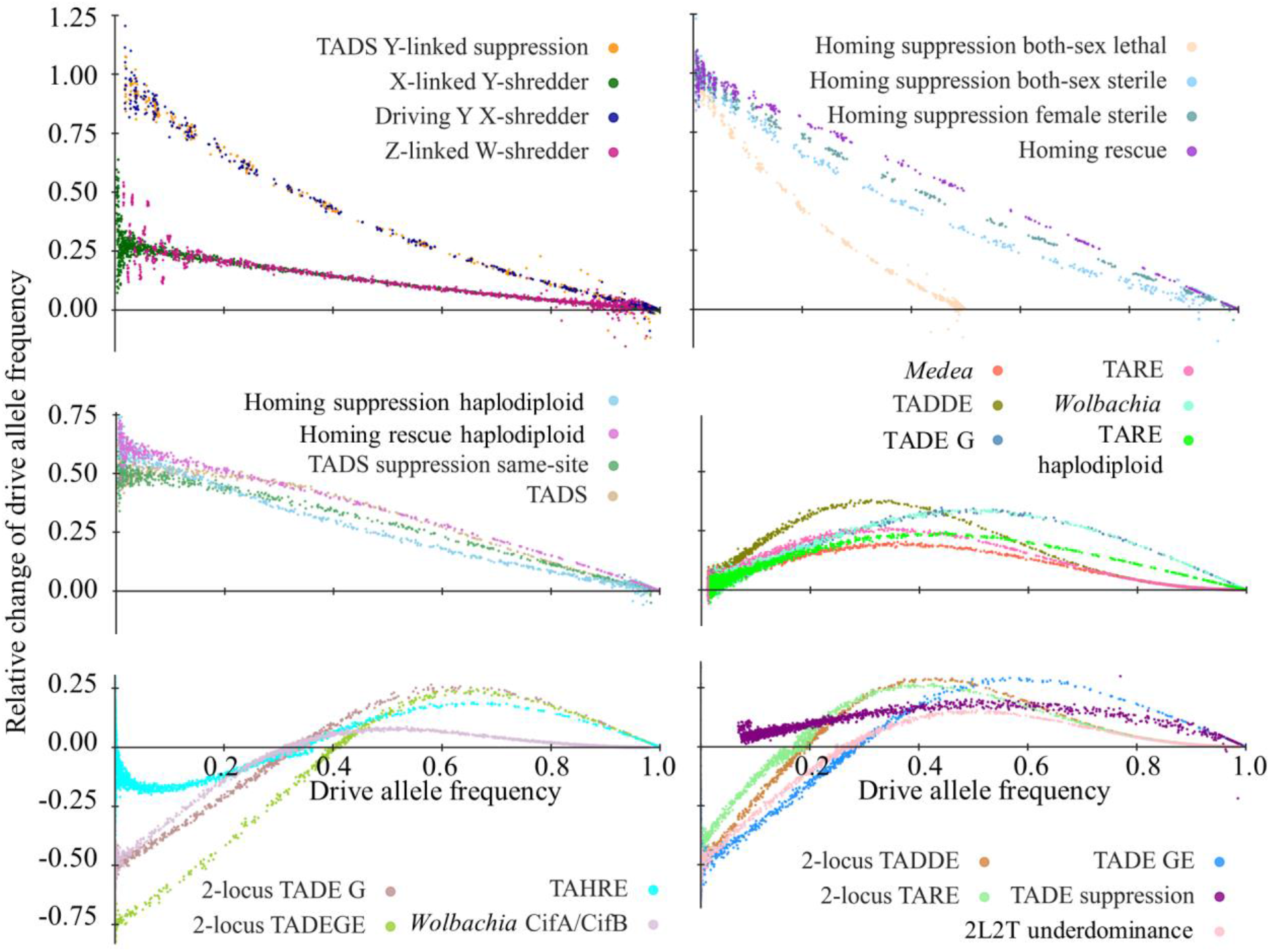
Drive performance dynamics in panmictic populations. Drive carriers were released into a panmictic population of wild-type individuals with several starting frequencies and ten replicates per starting frequency. The drive allele frequency was recorded for each generation. Simulations were stopped when the drive reached 0% or 100% frequency or when the population was eliminated. Each point’s position on the vertical axis represents the change in the number of wild-type alleles between two generations divided by the frequency in the earlier generation. The point’s value on the horizontal axis represents the total drive frequency in the earlier generation. Up to a few generations of data was removed from the beginning of each simulation so that displayed points would be unaffected by the starting conditions.

From this perspective, our gene drives can be clearly divided into the same six categories we saw in our wave speed and introduction threshold analysis in table 1. The Y-linked and homing drives can double in heterozygotes, and they thus have a relative increase rate of 1 at low frequencies (Figure 6). This clearly makes them pulled waves. Their rate of increase declines at higher frequencies, linearly for the modification drive and more rapidly at first for suppression drives. Both-sex lethal homing suppression drives cannot go above 50% frequency because wild-type alleles are required for viability. Next are the haplodiploid homing and TADS drives, which can increase rapidly, but only in one sex. At low drive frequency, their relative rate of increase is thus lower than 1 but it is still above zero, declining with increasing drive frequency. The X-linked and Z-linked shredder drives follow a similar pattern, but with an even lower starting level at low drive frequency because these drive alleles are less commonly found at any particular time in an individual where they can conduct shredding activity (thus increasing the drive frequency).

The frequency-dependent drives have a markedly different pattern (Figure 6). For the ones with zero threshold in ideal form, they have a relative increase of zero at low density. This makes it substantially more difficult for the leading edge of a wave to advance, though it is still possible. The middle part of the wave, though, likely makes a much greater contribution to wave advance. Yet, all of these drives (except for TADE suppression, as discussed previously) have a substantially higher rate of advance than the drives with thresholds in ideal form. These drives have a negative growth rate below their threshold, and they have the slowest wave speeds. All increases in drive frequency would thus come from deeper into the wave front, putting these solidly in the category of pushed waves.

### Relationship of drive performance in panmictic population and continuous space

Examining how a drive increases in frequency (Figure 6) can help with understanding the behavior of the drive in continuous space, but this method has some limitations. Some drives take a long time to increase to 100% even in ideal form (with the remainder often being nonfunctional resistance alleles), and this part of the wave tends to be far enough from the wave front that it has no effect on the wave’s propagation. Suppression drives may increase in frequency, but they also suppress the population, which has different dynamics. All drives, however, eliminate wild-type alleles fairly rapidly in ideal form. Such elimination could involve their removal from the population, or simply their conversion into drive alleles or disrupted alleles. This elimination is directly proportional to the wave advance speed in spatial models. Thus, examining this eliminate rate in panmictic models may provide a close proxy to a drive’s spatial performance.

We first collected data from the same panmictic simulations described in the previous section and displayed in Figure 6. However, instead of examining the relative drive rate of increase, we plotted the absolute change in wild-type allele count relative to the starting number in the whole population, plotted against the wild-type allele frequency in the earlier generation (Figure S6). This led to broadly similar patterns, but the advantage of zero-threshold drives at low drive frequency (high wild-type frequency) was obscured because even rapid drives will only remove wild-type alleles at a low absolute rate if the drive frequency is low. For the sex-linked drives that bias the sex ratio, removal of the opposing chromosome is an important part of drive dynamics and is relevant to overall drive advance. We thus counting opposing chromosomes as wild-type alleles and plotted the same data (Figure S7). For the Y-linked drives, wild-type X chromosomes would always be present with drive alleles, so the wild-type frequency can never fall below 0.5.

To examine how panmictic removal of wild-type alleles was related to drive wave speed (which must be proportional to the removal of wild-type alleles in our 1-dimensional model), we calculated the integral of curves in Figure S6 for most drives, using Figure S7 for the sex-linked drives that bias the sex ratio. This was plotted against the wave speed for each type of drive (Figure 7). In general, these are well correlated, with more negative integrals of wild-type allele change corresponding to faster drive wave speeds. Indeed, if the wave shape was completely linear with respect to wild-type frequency at the wave front, we would expect them to have a perfectly linear correlation.

**Figure 7.**
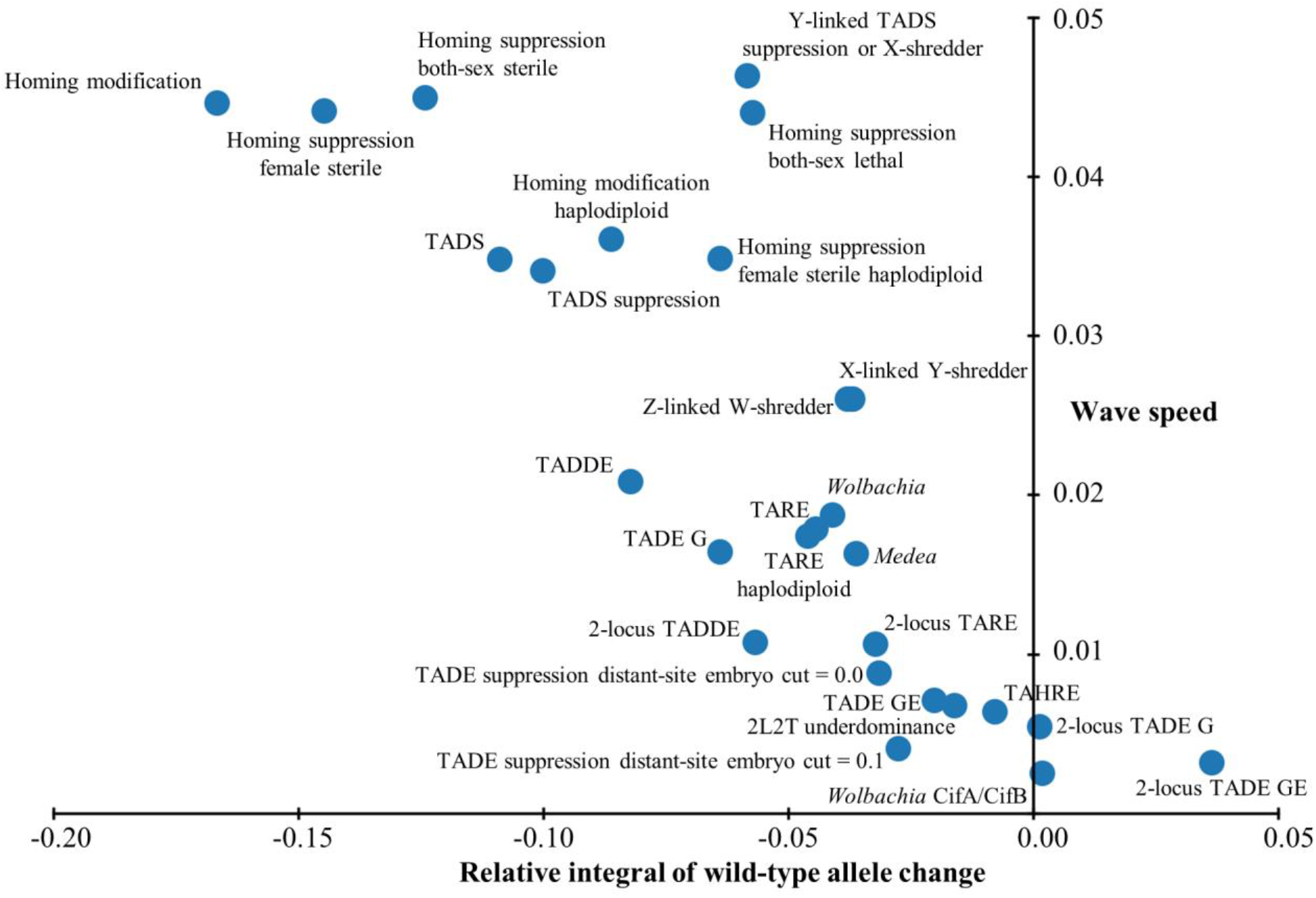
Relationship between drive wave speed and panmictic performance. For ideal drives, we obtained the integral of relative wild-type allele removal in the presence of the drive from 0 to 1 wild-type frequency (see Figures S6-S7 for data used for these integrals). This was plotted against the ideal drive wave advance speed.

However, some drives deviate from this hypothetical pattern. One prominent exception is the Y-linked suppression drives, even after adjusting for removal of wild-type X-chromosomes. This is perhaps because their adjusted integral from Figure 7 only had half the width as our other integrals, giving them an unfairly low value. Yet, when considering their drive wave, there would be no reason to have a different width than other drives, so perhaps their “effective integral” could be considered as double the plotted value, putting them more in-line with other points. This same phenomenon applies to the both-sex lethal homing suppression drive.

The most obvious deviation from a linear correlation are the three drives that has a positive integral, the 2-locus TADE variants and the *Wolbachia* CifA/CifB drive. This occurs because they have high introduction thresholds, and drive alleles are rapidly eliminated at low frequencies. However, if wild-type alleles were increasing in frequency on average across the drive wave, then the drive wave would be receding instead of advancing. These drives certainly have low wave speed, but they can clearly form an advancing wave. To explain this, these drives must have a wider wave where wild-type frequency is low to medium, thus making these regions more important for overall wave speed than the narrow area in the front of the wave where drive alleles are being removed. It is these sorts of deviations from a linear wave shape that can explain the rest of the discrepancies between our data and a perfectly linear correlation. In general, faster removal of wild-type (or increase in wild-type) would likely represent a narrower fraction of the wave compared to areas of slower change, thus contributing less to wave speed than its magnitude in Figure S6 would indicate.

## Discussion

In this study, we evaluated a large number of different gene drives using a unified framework, facilitating cross-drive comparisons. We examined the form of advancing drive waves and the speed of such waves as a function of germline drive efficiency, embryo cut rates (for CRISPR-based drives), drive fitness costs, low-density growth rate, and dispersal rates. We compared this wave advance speed with drive performance in panmictic populations and found some interesting differences.

As can be seen in our integral results of panmictic wild-type allele removal, the performance of drives in panmictic populations is well-correlated with spatial wave advance speed, but there are several important differences, particularly for pushed waves. While the “front” part of a drive wave where wild-type frequency is high is sufficient to pull zero-threshold drives rapidly with dispersal, regardless of what is happening in the back of the wave, the middle part of frequency-dependent drives is essential for moving the drive forward. This changes the shape of the drive wave, which can give more prominence to regions of the space where the drive is at higher frequency. A slow drive can have a wider wave, giving more interaction between drive and wild-type and thus allowing it to maintain higher speed than one would expect when considering only panmictic drive performance. For drives with high thresholds, the front part of the wave involves faster elimination of drive alleles than wild-type alleles, and the drive is only sustained by migration from further back. In this case, the shape of the wave matters even more, with some drives even having positive wild-type integrals if each part of the entire wild-type frequency spectrum is considered with equal weight, highlighting the difference between panmictic and spatial drive spread characteristics.

While homing drives are among the fastest, they are also the most sensitive to imperfect drive performance parameters, rapidly losing speed as both germline efficiency and embryo cut rates decline. This is because resistance alleles, even when they can be removed, also stop drive conversion, which is the optimal mechanism by which a drive can increase its frequency. TADE drives also tend to be greatly slowed by cleavage in the early embryo because this often results in removal of drive alleles^70^. TADE drives thus gain thresholds with even modest levels of embryo reistance^25,102^. For other drives, such as those based on TARE and 2-locus systems, drive performance has a reduced effect on drive speed.

One critical question for considering the deployment of a gene drive is whether it can form a self-sustaining wave of advance in the first place. In our analysis, we saw that poor performance parameters for suppression drives can result in drive failure. However, if the germline efficiency is low, then the drive might lack enough genetic load to substantially reduce or eliminate the population in the first place. However, both-sex homing drives, haplodiploid homing drives, and TADE suppression drives are all vulnerable to high embryo cut rates even if germline performance is adequate. TADE suppression drive is particularly vulnerable to embryo cutting because it is a frequency dependent drive that quickly gains a high introduction threshold, yet it is also a suppression drive that reduces its own density^70^. This phenomenon has been explored mathematically for other suppression drives^57,103^. When fitness costs are considered, both modification drives and suppression drives can rapidly lose their ability to spread, particularly for frequency-dependent drives with high thresholds. This tends to occur even before a 50% introduction threshold is released. Thus, if a cargo gene has a moderate fitness cost, that might change the optimal drive for a particular scenario, and for high fitness costs, a tethered drive system may be necessary to spread the cargo if the drive is required to be confined to only a specific target population^24,37^.

One notable result from our study was that the low-density growth rate did not have a large effect on drive wave speed. As this parameter increases to approximately 3-5, many drives experienced a small increase in wave speed, and waves become viable for some weak drives even in the presence of fitness costs, but past this level, simulations show very little additional effect of further increases. This was somewhat unexpected because many drives influence the population level, even if they are not suppression drives, due to toxin-antidote effects. Stronger density dependence would allow such drives to maintain a higher population level at the interface with wild-type individuals and thus put more pressure on the wild-type population by migration, resulting in a faster drive wave. However, we used a fixed density-dependent curve (Beverton–Holt model) that does not provide a major fitness advantage until substantial reductions in density are observed. This was to avoid unstable populations that could grow substantially above carrying capacity in our discrete generation model. A different density-dependent competition curve^104^ could substantially change this results for some frequency-dependent drives such as TADE suppression.

One possible application of the drive wave speed would be to determine an optimal release pattern for a given gene drive. A fast drive would require a smaller resource investment to cover the same area in a given time interval, potentially allowing wider coverage, even if wider dispersion is costly. Slower drives would require multiple releases in different locations to achieve the same drive coverage for the same limited time interval. However, such slower drives also require higher critical release sizes^59,61,70,105,106^, which can potentially make this difficult and could perhaps lead to an optimal strategy of focusing on one limited region at a time, or planning for a strategy that takes longer to fully mature. In any case, an expected wave speed of a drive could allow for optimization of a monitoring program, potentially reducing costs of any study involving gene drive releases.

Due to our desire to compare drive performance in a general, broadly applicable framework, our model is necessarily simplified compared to real populations. In particular, we used discrete generations, which is not representative of most species, and considered a one-dimensional arena with no spatial heterogeneity. Complex lifecycle characteristics that are typical of many species^65,66^ were excluded, and competition was also only considered between adults. It was also restricted to affecting female fecundity. While representative of some species, other important species such as mosquitoes tend to produce a lot of juveniles, which then compete intensely for resources that they need to develop into adults, thus affecting viability. Other ecological characteristics were also ignored, such as seasonality and the exact nature of the density-dependence function^104^, which could vary between species. This in particular could substantially affect the speed of drives that suppress a population and are frequency-dependent (both suppression drives and modification drives that reduce the population density at the wave front). These issues in our model mean that we are limited when making quantitative predictions for any specific species. Though comparative drive performance would likely be similar for a given species in many situations, exact timing and speed intervals would be greatly affected by ecological characteristics, in particular migration/dispersal rates.

In summary, we measured the wave speed of many types of gene drives with varying drive performance and ecological parameters, revealing several interesting differences in relative wave speed between drives and determining when drive waves become nonviable. These results can facilitate consideration of optimal drive types for various situations and also inform drive release patterns if such drives are deployed in the future. Critical performance parameters are identified for some drives, which could allow for more focused research for improvement of these performance characteristics. Our results also provide a point of comparison for drive performance measures in both panmictic populations and continuous space populations, showcasing the importance of evaluating even simpler modification drives in spatial frameworks.

## Acknowledgements

Thanks to the High-Performance Computing Platform of the Center for Life Science at Peking University for assistance with cluster-based data collection. This study was supported by laboratory startup funds from Peking University and the NSFC Overseas Youth Fund.

## Supplemental Information

**Table S1.** List of gene drives not assessed in this study.

| Drive | Type | Threshold | Speed | Activity | Status |
| --- | --- | --- | --- | --- | --- |
| Haploinsufficient underdominance | M | 0.5 | N/A | Any | Flies<sup>28,29</sup> |
| TADE GES | M | 0.5 | N/A | GES | Theoretical<sup>+</sup><sup>102</sup> |
| Reciprocal chromosome translocations | M | 0.5 | N/A | None | Flies<sup>107</sup>, Mosquitoes<sup>108,109</sup> |
| 1-locus 2-drive TARE | M | 0.61 | N/A | Any | Theoretical<sup>+</sup><sup>102</sup> |
| 1L2T underdominance | M | 0.67 | N/A | Any? | Theoretical<sup>33</sup> |
| Compound chromosome translocations | M | 0.8 | N/A | None | Flies<sup>107</sup> |
| 2-locus TADE suppression | S | 0.83/0.88 | N/A | G/GE | Theoretical<sup>+</sup><sup>102</sup> |
| TAHRE suppression | S | 0.96 | N/A | GE | Theoretical<sup>102</sup> |

Drive type: M = modification, S = suppression
Threshold refers to the introduction threshold below which the drive will be eliminated. Released individuals are homozygotes for modification drives and heterozygotes for suppression drives.
N/A = drive cannot form a wave of advance
Activity refers to the period in which engineered drive mechanism components are active. G = germline, E = early embryo, S = somatic cells
+could likely be engineered with demonstrated components and existing knowledge

**Figure S1.**
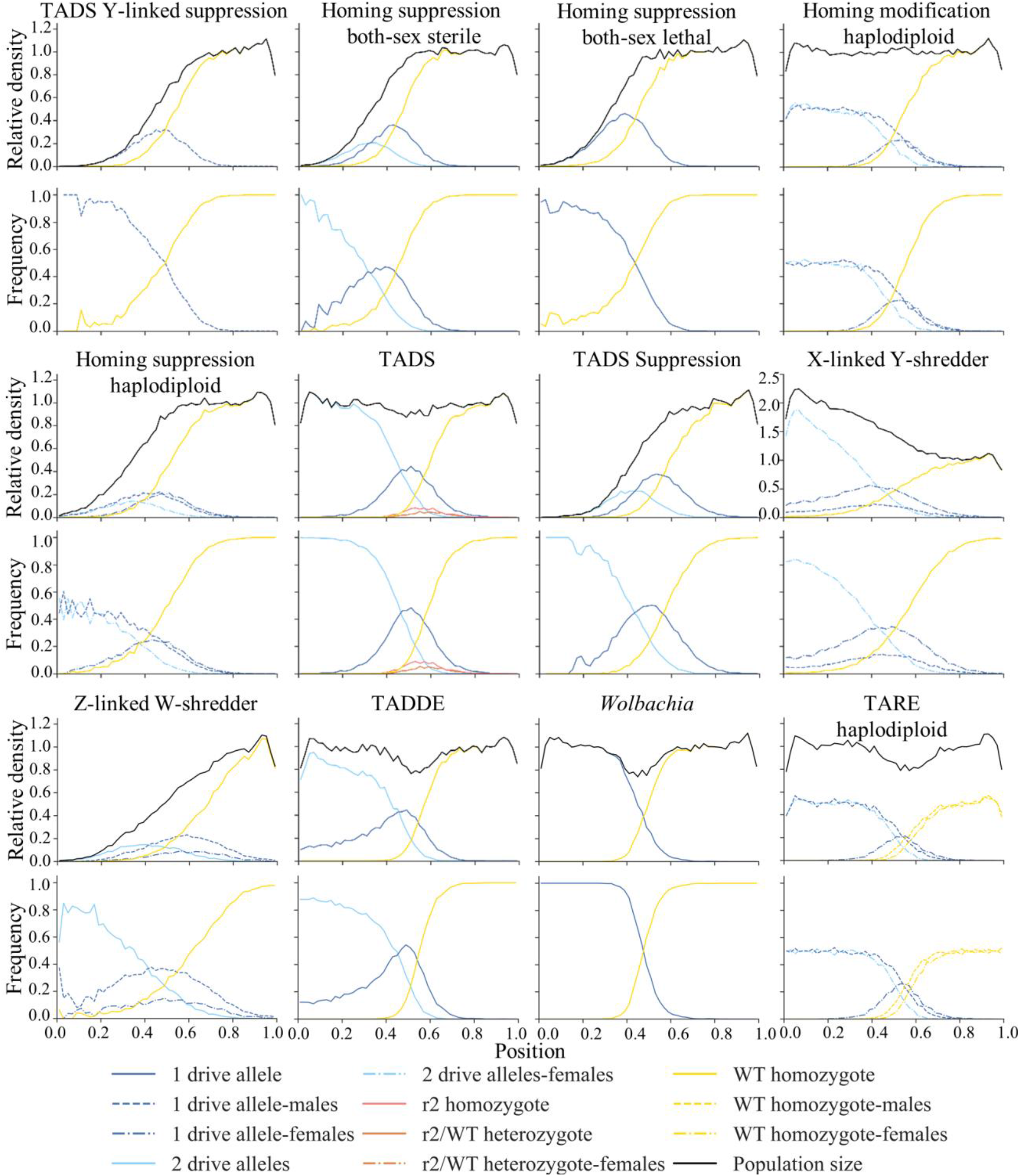
Snapshots (first half). The relative density (compared to wild-type in the absence of drive) of several genotype categories is displayed for an advancing drive wave, as well as the total population density. The lower panels show the frequencies of each genotype category. Due to space limitations, this figure is broken up into two panels. Note that the density scale is different from the X-linked Y-shredder. The graphs are divided into 50 points covering the whole one-dimensional space of length 1.

**Figure S1.**
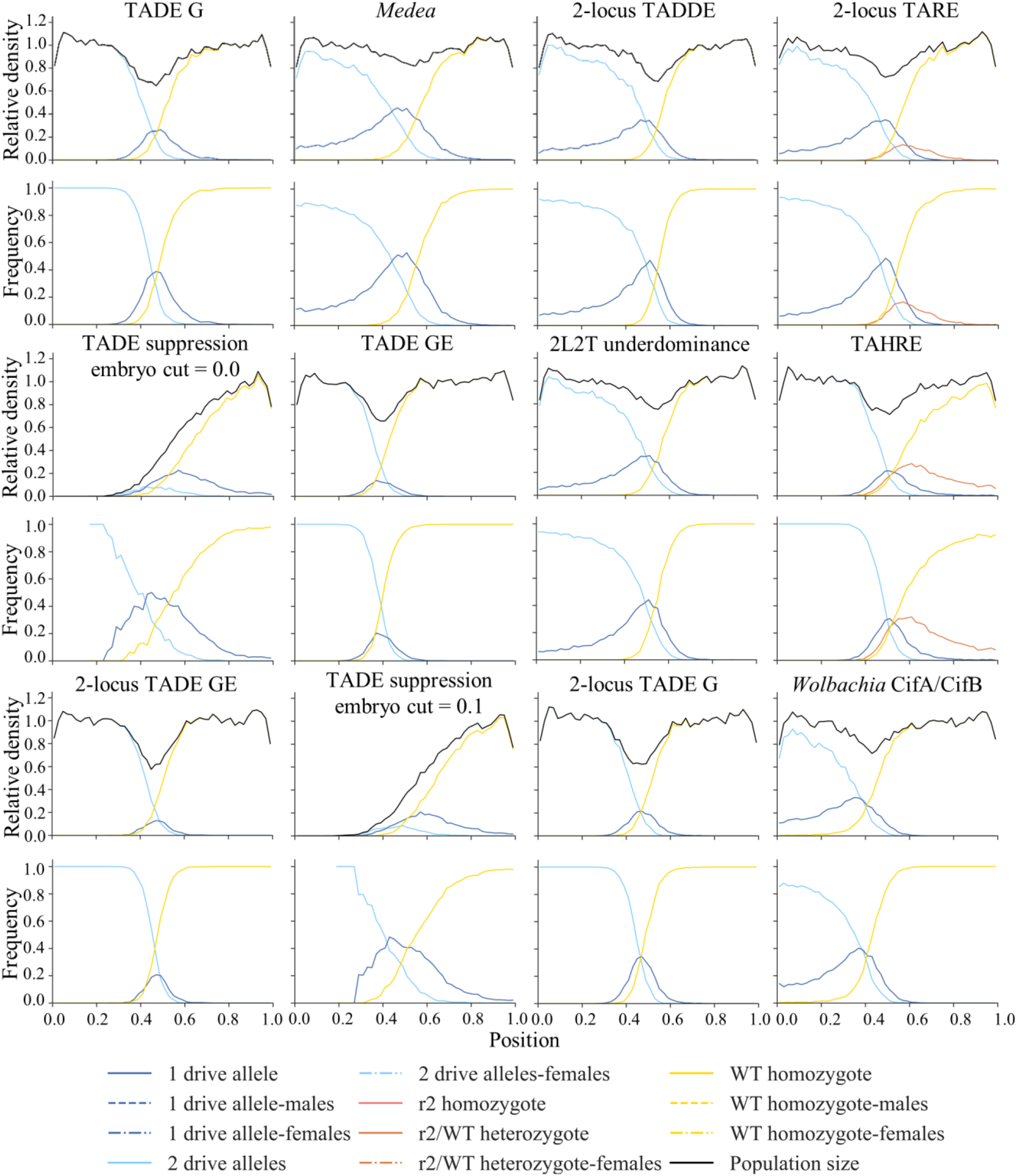
Snapshots (second half). The relative density (compared to wild-type in the absence of drive) of several genotype categories is displayed for an advancing drive wave, as well as the total population density. The lower panels show the frequencies of each genotype category. Due to space limitations, this figure is broken up into two panels. The graphs are divided into 50 points covering the whole one-dimensional space of length 1.

**Figure S2.**
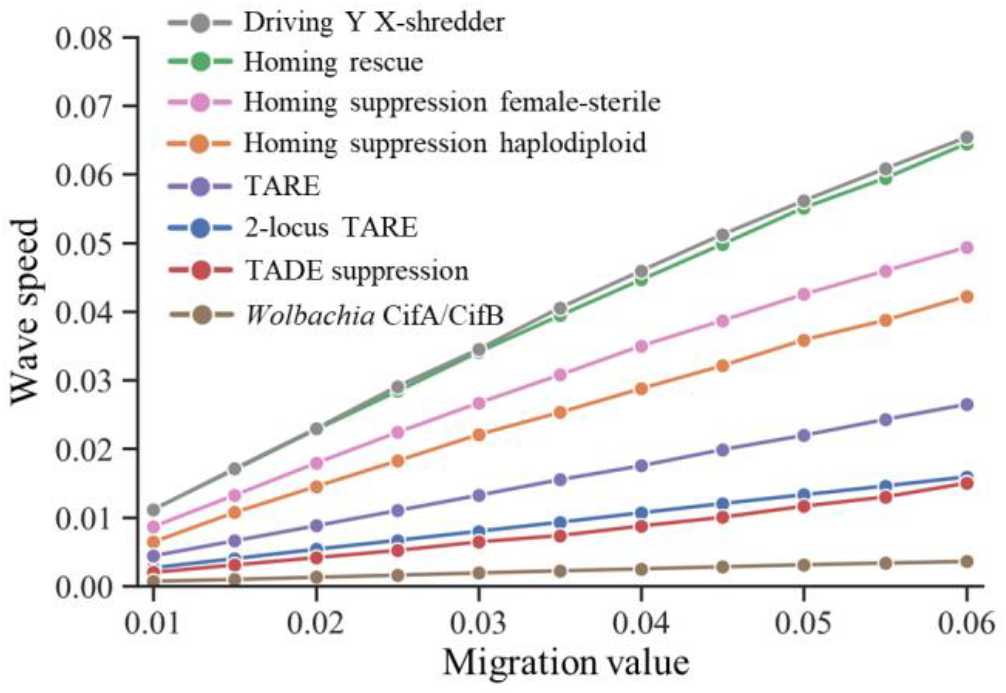
Effects of migration value on drive wave speed. Drive carriers were released into the left edge of a population of wild-type individuals, and the wave speed was measured for varying migration value. Each point represents at least 200 simulations.

**Figure S3.**
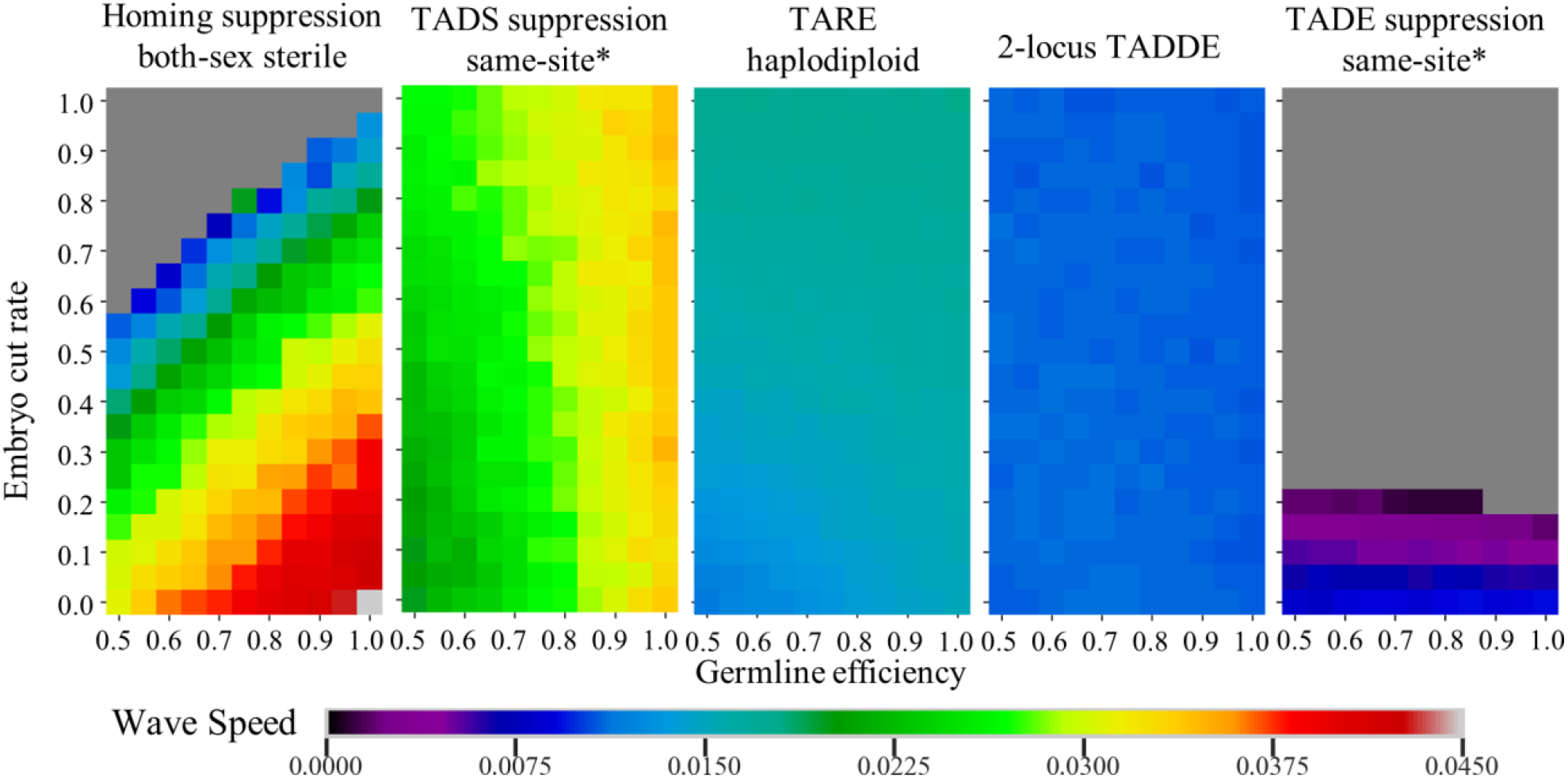
Additional heatmaps showing the effect of germline efficiency and embryo cut rate on wave speed. Drive carriers were released into the left edge of a population of wild-type individuals, and the wave speed was measured for varying germline efficiency and embryo cut rate. Each point represents the average of 20 simulations. Grey means that a wave of advance could not form. * means that somatic fitness costs were used instead of multiplicative fitness.

**Figure S4.**
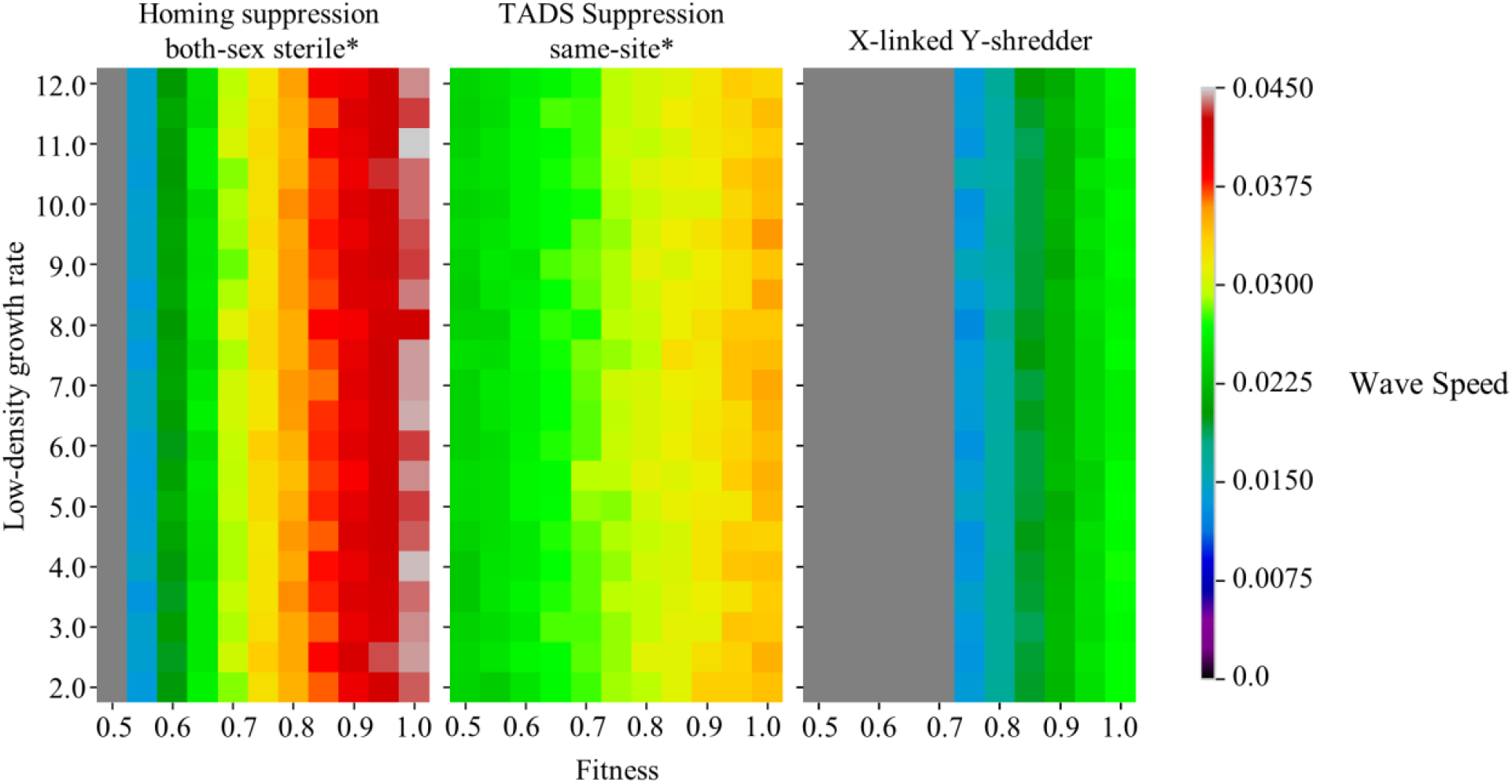
Additional heatmaps showing the effect of low-density growth rate and fitness value. Drive carriers were released into the left edge of a population of wild-type individuals, and the wave speed was measured for varying fitness and low-density growth rate. Each point represents the average of 20 simulations. Grey means that a wave of advance could not form. * means that somatic fitness costs were used instead of multiplicative fitness.

**Figure S5.**
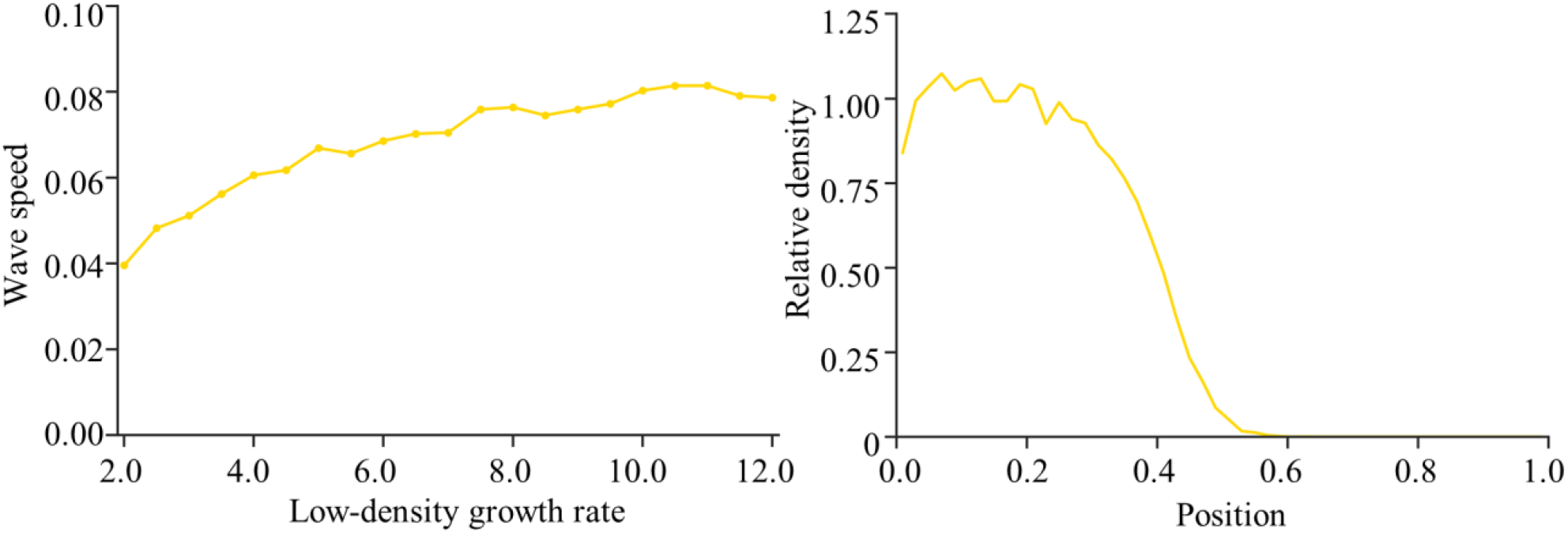
Wild-type waves of advance into empty space. 1,000 wild-type individuals were released into the left edge of an empty space with a capacity of 10,000, and the wave speed was measured for varying low-density growth rate (left panel). Each point represents the average of 20 simulations. The right panel shows a representative example of the density as wild-type individuals advanced into empty space.

**Figure S6.**
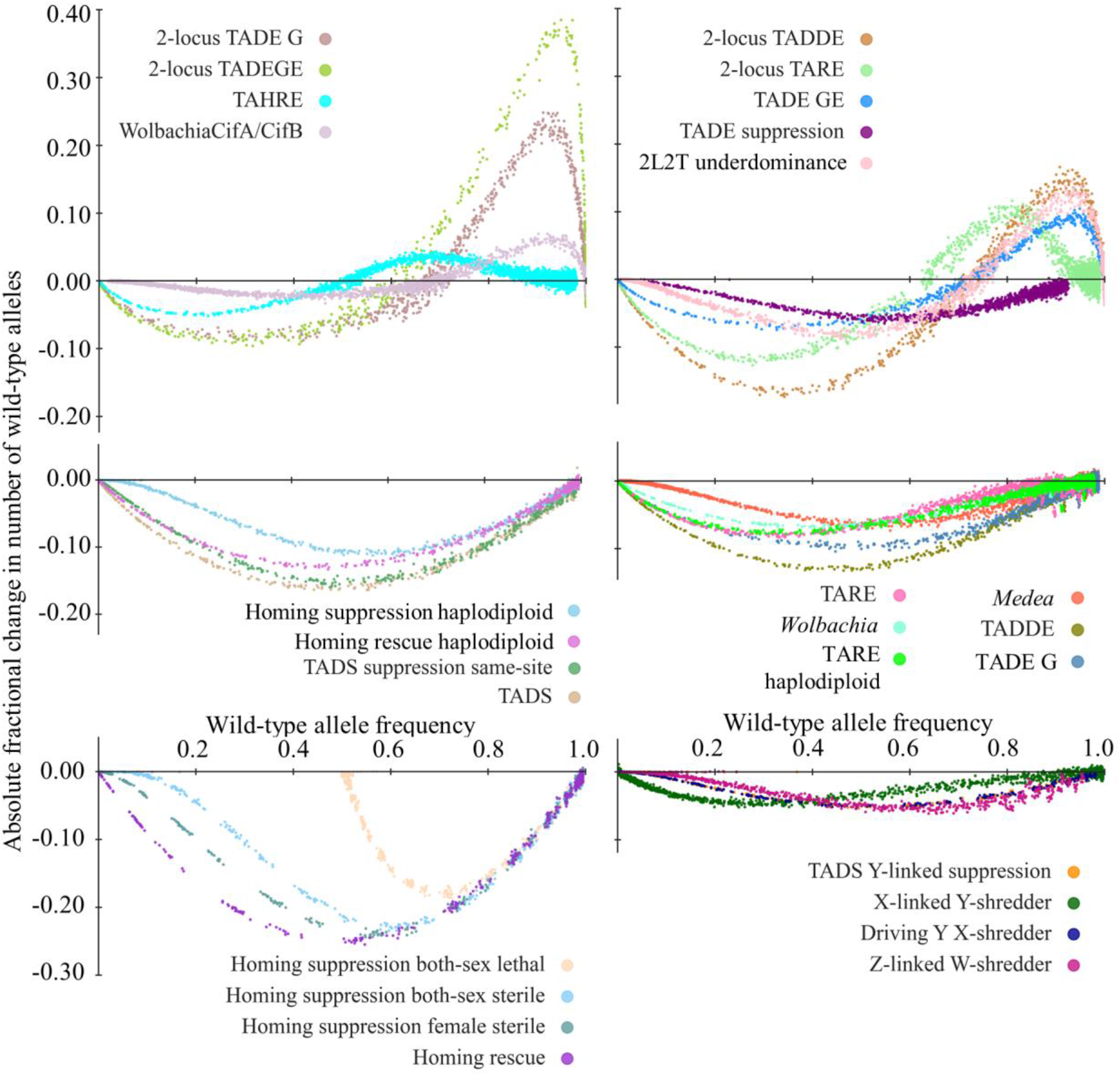
Absolute fractional change of the number of the wild-type allele. Drive carriers were released into a panmictic population of wild-type individuals with several starting frequencies and ten replicates per starting frequency. The wild-type allele frequency was recorded for each generation. Simulations were stopped when the drive reached 0% or 100% frequency or when the population was eliminated. Each point’s position on the vertical axis represents the change in the number of wild-type alleles between two generations divided by the total number of wild-type alleles in the population at equilibrium without any drive (200,000). The point’s value on the horizontal axis represents the wild-type frequency in the earlier generation. Up to a few generations of data was removed from the beginning of each simulation so that displayed points would be unaffected by the starting conditions.

**Figure S7.**
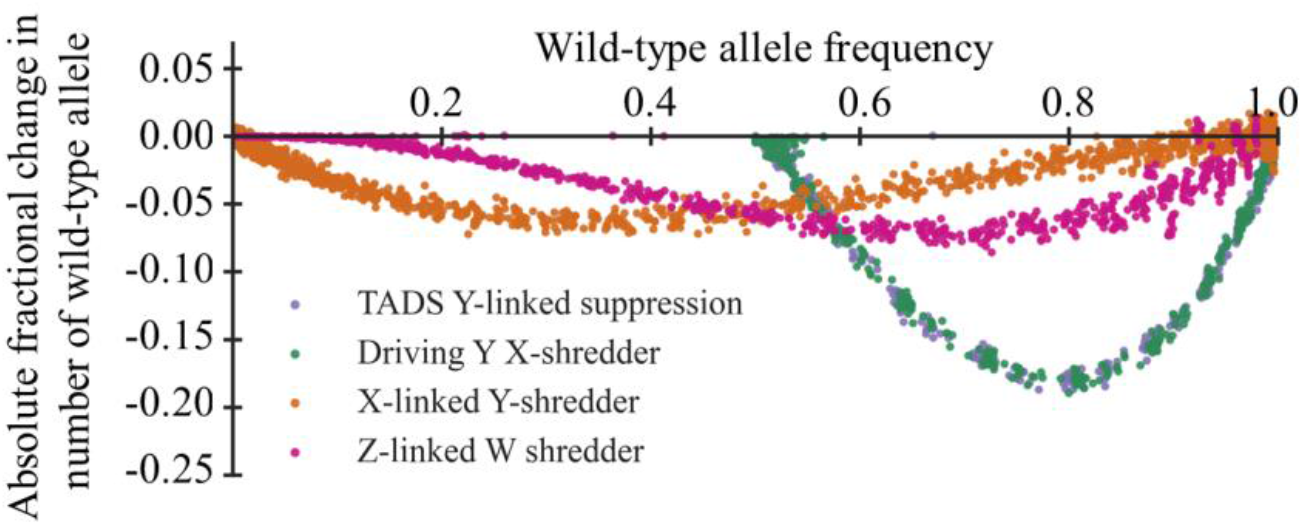
Absolute fractional change of number of wild-type allele. This shows the same data as in Figure S6 for sex-linked suppression drives, except the wild-type alleles were considered to include the sex chromosome that the drive was not on, in addition to wild-type variants of the drive’s chromosome.

